# Redox Nanomedicine Cures Chronic Kidney Disease (CKD) by Mitochondrial Reconditioning

**DOI:** 10.1101/2021.03.14.435287

**Authors:** Aniruddha Adhikari, Susmita Mondal, Tanima Chatterjee, Monojit Das, Pritam Biswas, Soumendra Darbar, Hussain Alessa, Jalal T. Al-Thakafy, Ali Sayqal, Saleh A. Ahmed, Anjan Kumar Das, Maitree Bhattacharyya, Samir Kumar Pal

**Affiliations:** Department of Chemical, Biological and Macromolecular Sciences, S. N. Bose National Centre for Basic Sciences, Block JD, Sector 3, Salt Lake, Kolkata-700106, India; Department of Biochemistry, University of Calcutta 35, Ballygunge Circular Road, Kolkata-700019, India; Department of Zoology, Uluberia College, University of Calcutta, Uluberia, Howrah-711315, India; Department of Zoology, Vidyasagar University, Rangamati, Midnapore-721102, India; Department of Microbiology, St. Xavier’s College, 30, Mother Teresa Sarani, Kolkata-700016, India; Research & Development Division, Dey’s Medical Stores (Mfg.) Ltd, 62, Bondel Road, Ballygunge, Kolkata-700019, India; Department of Chemistry, Faculty of Applied Sciences, Umm Al-Qura University, 21955 Makkah, Saudi Arabia; Chemistry Department, Faculty of Science, Assiut University, 71516 Assiut, Egypt; Department of Pathology, Calcutta National Medical College and Hospital, 32, Gorachand Rd, Beniapukur, Kolkata-700014, India

**Keywords:** Nanomedicine, Redox healing, ROS balancing, Renal fibrosis, Nephrotoxicity, Mitochondrial reconditioning

## Abstract

Targeting reactive oxygen species (ROS) while maintaining cellular redox signaling is crucial in the development of redox medicine for the therapeutic benefit as the origin of several prevailing diseases including chronic kidney disease (CKD) is linked to ROS imbalance and associated mitochondrial dysfunction. Here, we have shown that an indigenously developed nanomedicine comprising of Mn_3_O_4_ nanoparticles duly functionalized by biocompatible ligand citrate (C-Mn_3_O_4_ NPs) can maintain cellular redox balance in an animal model. We developed a cisplatin-induced CKD model in C57BL/6j mice where severe mitochondrial dysfunction resulting in oxidative distress lead to the pathogenesis. Four weeks of treatment with C-Mn_3_O_4_ NPs restored renal function, preserved normal kidney architecture, ameliorated overexpression of pro-inflammatory cytokines, and arrested glomerulosclerosis and interstitial fibrosis in CKD mice. A detailed study involving human embryonic kidney (HEK 293) cells and isolated mitochondria from experimental animals revealed that the molecular mechanism behind the pharmacological action of the nanomedicine involves protection of structural and functional integrity of mitochondria from oxidative damage, the subsequent reduction in intracellular ROS, and maintenance of cellular redox homeostasis. To the best of our knowledge, such studies that efficiently treated a multifaceted disease like CKD using a biocompatible redox nanomedicine are sparse in the literature. Successful clinical translation of this nanomedicine may open a new avenue in redox-mediated therapeutics of several other diseases (e.g., diabetic nephropathy, neurodegeneration, and cardiovascular disease) where oxidative distress plays a central role in pathogenesis.

## INTRODUCTION

Reactive oxygen species (ROS) have long been considered as an unwanted but inevitable byproduct of aerobic oxygen metabolism [1]. Excessive generation of ROS may lead to tissue damage and numerous undesired physiological consequences. Increased ROS level is linked to inflammation, aging, and pathogenesis of diseases like diabetes, cancer, atherosclerosis, chronic kidney disease (CKD), and neurodegeneration [2-5]. Recent understanding about the pivotal role of ROS as secondary messengers in cellular signaling to control processes like metabolism, energetics, cell survival, and death lead to a paradigm shift to the traditional “oxidants are bad – antioxidants are good” based simplistic view of redox biology [6-10]. Apathy towards the paradox between lethality of excessive intracellular ROS (oxidative distress) and beneficial role of low concentration ROS (oxidative eustress) is the major underlying reason behind the failure of conventional antioxidant therapies using natural or synthetic antioxidants (e.g., α-tocopherol, ascorbic acid, β-carotene, curcumin, and numerous polyphenols present in the diet) that along with scavenging intracellular free radicals in a stoichiometric way, insulates redox signaling [10-12]. Moreover, meta-analyses of clinical trials show that conventional antioxidants are not only ineffective, but harmful, and even increase mortality [12, 13]. The understanding that proper cell functioning critically requires a dynamic balance between oxidative eustress and distress (i.e., cellular redox homeostasis) forms the conceptual framework of redox medicine, a novel therapeutics that passivates the oxidative distress while maintains the normal redox circuitry [10, 12, 14-16]. The cellular redox dynamics and its regulations, however, are still largely elusive because of the lack of effective pharmacological interventions [17]. In this regard, biocompatible transition metal oxide nanoparticles with potential electron-donating as well as accepting capability could be a viable option provided they are stable in the biological system, able to assimilate in the targeted tissue, and functional in the physiological *milieu*.

Recently, we have shown that spinel structured citrate functionalized Mn_3_O_4_ nanoparticles (C-Mn_3_O_4_ NPs) have the unique ability to generate ROS in dark, and when injected into jaundiced animals can selectively degrade bilirubin (a toxic byproduct of heme metabolism) without showing adverse effects to other blood parameters [18]. In the *in vitro* reaction system, we found that the nanoparticles can catalytically scavenge free radicals particularly H_2_O_2_. The microenvironment-controlled (i.e., presence of ROS, subsequent changes in pH and dissolved O_2_) dynamic equilibrium between disproportionation and comproportionation involving Mn^3+^, Mn^4+^ and Mn^2+^ charge states present in the hausmannite structure of C-Mn_3_O_4_ NPs is responsible for such dual activity [19-21]. Hence, depending upon the intracellular redox condition, the nanoparticle has the potential to balance the oxidative distress and eustress, the most important feature of a redox medicine.

In this study, our major aim was to evaluate the potential of C-Mn_3_O_4_ NPs as a redox medicine against CKD. CKD, the progressive decline in kidney function, is one of the most serious global public health problem (with 8-16% worldwide prevalence) that originates from redox imbalance due to mitochondrial dysfunction and have no effective medicine till date [22-26]. To evaluate the therapeutic potential of C-Mn_3_O_4_ NPs we used a cisplatin-induced C57BL/6j mice model of CKD. The mechanistic details of their pharmacological action in the maintenance of redox homeostasis and mitoprotection were further explored using cellular (human embryonic kidney cell, HEK 293) as well as the animal model.

## RESULTS

### Designing aqueous soluble C-Mn_3_O_4_ NPs to target kidney cells

The size, surface charge, and surface functionalized ligands determine the biodistribution of a nanomaterial inside living organisms. Previous studies have reported that particles with less than 8 nm diameters having moderate to high surface negative charge tend to accrue in the renal system [27]. So, care was taken at the time of synthesis to control the size of the Mn_3_O_4_ nanoparticles within the range of 6 nm. The transmission electron micrograph (TEM) of C-Mn_3_O_4_ NPs shows monomodal distribution of nearly spherical particles with an average diameter of 5.58±2.42 nm (Figure 1a). High resolution (HR) TEM image of a single nanoparticle confirms the crystalline nature with clear atomic lattice fringe spacing of 0.311±0.02 nm (Figure 1a-inset) corresponding to the separation between (112) lattice planes of hausmannite Mn_3_O_4_ crystal. All x-ray diffraction (XRD) peaks corresponding to (101), (112), (200), (103), (211), (004), (220), (204), (105), (312), (303), (321), (224), and (400) planes of C-Mn_3_O_4_ NPs (Figure 1b) exactly reflected the tetragonal hausmannite structure of Mn_3_O_4_ with a lattice constant of a=5.76Å and c=9.47Å and space group of I41/amd described in the literature (JCPDS No. 24-0734). The absence of any additional peak from other phases indicates the high purity of the synthesized material.

**Figure 1.**
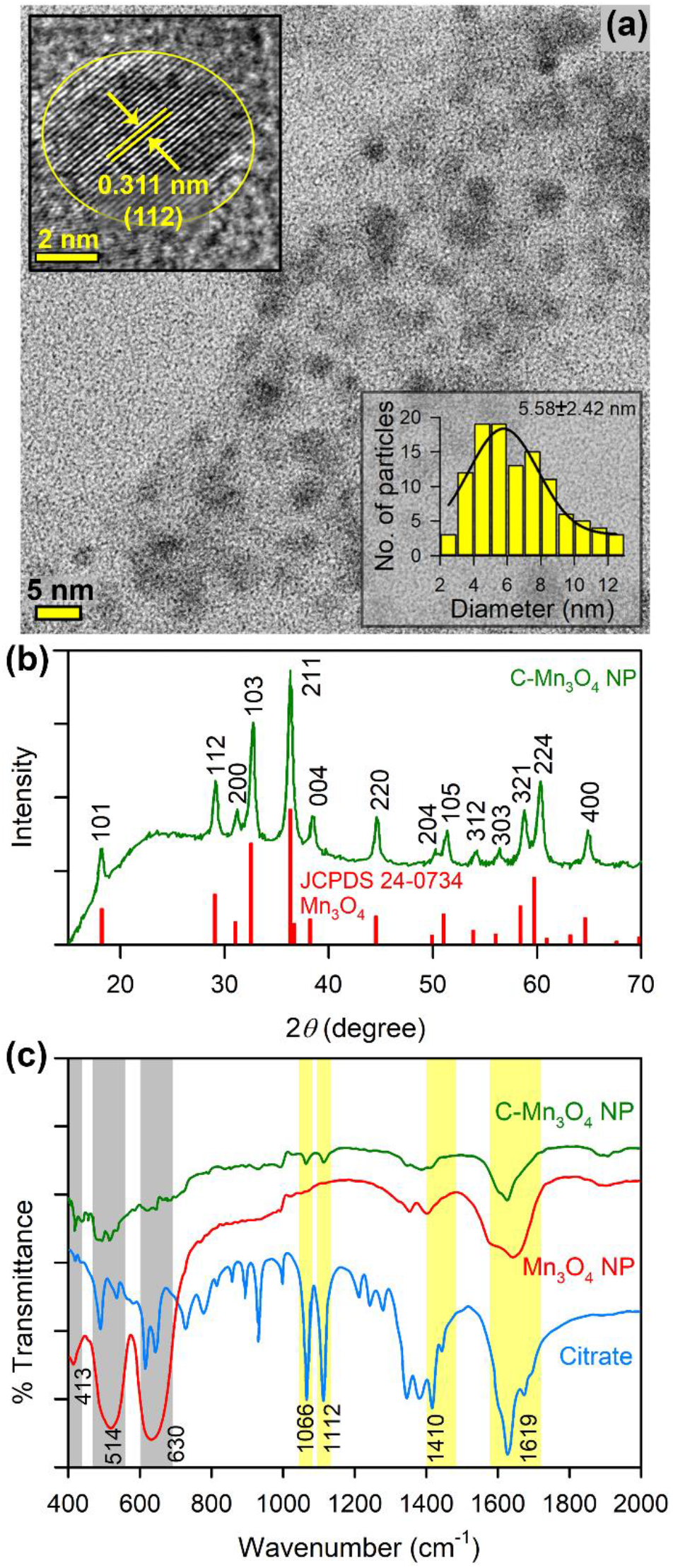
Characterization of C-Mn_3_O_4_ NPs. (a) TEM image of C-Mn_3_O_4_ NPs shows spherical shape of the nanoparticles with monomodal distribution. Inset shows HRTEM image of single nanoparticle having high crystalline structure with 0.311 nm interfringe distance corresponding to the (112) plane. The other inset shows histogram of size distribution of the particles with average diameter of 5.58±2.42 nm. (b) Experimental XRD peaks of the nanoparticle exactly matches to that of literature defined one for Mn_3_O_4_ hausmannite (JCPDS No. 24-0734). (c) FTIR spectra of C-Mn_3_O_4_ NPs, Mn_3_O_4_ NPs, and citrate. Perturbation at Mn-O stretching at 413, 514, 630 cm^-1^ (shaded grey) of Mn_3_O_4_ NPs and carboxylic groups at 1066, 1112, 1410, 1619 cm^-1^ (shaded yellow) of citrate confirms strong covalent binding between citrate and the nanoparticle.

Surface functionalization with carboxyl rich ligand tri-sodium citrate not only made the nanoparticles biocompatible and aqueous soluble but also helped the surface charge to be negative (i.e., zeta potential, ξ=-12.23±0.6 mV with electrophoretic mobility -0.96±0.05 μcmV^-1^s). Furrier transformed infrared (FTIR) spectra were used to confirm the binding of citrate to the surface of the nanomaterial (Figure 1c). Broadening of the 630, 514 and 413 cm^-1^ bands associated with stretching vibrations of Mn-O and Mn-O-Mn bonds of Mn_3_O_4_ NPs along with substantial disruption of both symmetric (1410 cm^-1^) and asymmetric (1619 cm^-1^) stretching modes of carboxylates (COO^-^) of citrate indicates a strong covalent interaction between them.

Previously we showed that C-Mn_3_O_4_ NPs can selectively degrade bilirubin without affecting other blood parameters [27]. Here, initially, we evaluated their potential to scavenge H_2_O_2_ in an *in vitro* system using Rose Bengal (RB) degradation assay. RB has a distinct absorption peak at 540 nm. In the presence of H_2_O_2_, RB degenerates with a subsequent decrease in the 540 nm absorbance. When added to the reaction mixture, C-Mn_3_O_4_ NPs efficiently prevented the RB from H_2_O_2_ mediated degradation (Supplementary Figure S1) indicating its strong radical scavenging potential towards H_2_O_2_.

### C-Mn_3_O_4_ NPs maintain redox balance in HEK 293 cells against H_2_O_2_-induced oxidative distress

To test the ability of C-Mn_3_O_4_ NPs to combat oxidative stress in the cellular environment, we used a cell-based approach. The HEK 293 cells pretreated with different concentrations of nanoparticles (3.75 to 60 μg mL^-1^) were exogenously exposed to H_2_O_2_ (1 mM) and cell viability was estimated using well known 2-(4,5-dimethylthiazol-2-yl)-2,5-diphenyltetrazolium bromide (MTT) assay (Figure 2a). The survival rate for H_2_O_2_ treated cells was ∼35% (p<0.001 compared to control). The C-Mn_3_O_4_ NPs protected the cells from H_2_O_2_ induced cell death in a dose-dependent manner. Cell viability reached a maximum of ∼85% and ∼88% (p<0.001 compared to H_2_O_2_ treated cells) in H_2_O_2_ exposed cells when pretreated with 30 and 60 μg mL^-1^ NPs respectively. Pretreatment of the cells with similar concentrations of the NPs alone did not cause significant cellular mortality. Therefore, we selected the 30 μg mL^-1^ concentration of C-Mn_3_O_4_ NPs for further experiments. Identical results were observed in the lactate dehydrogenase (LDH) assay (Figure 2b). The presence of a high concentration of H_2_O_2_ inside the cell caused oxidative damage to the plasma membrane resulting in an increased release of LDH, a cytosolic enzyme, into the surrounding cell culture medium. Pretreatment with C-Mn_3_O_4_ NPs protected the cells from H_2_O_2_ induced oxidative damage resulting in a ∼40% reduction in the LDH release (p<0.001 compared to H_2_O_2_ treated cells). To evaluate the scavenging of H_2_O_2_ by C-Mn_3_O_4_ NPs under stress condition, we monitored the intracellular oxidative stress using a ROS-sensitive fluorescence probe, dihydrodichloro-fluoresceindiacetate (DCFDA-H2). DCFDA-H2 is transported across the cell membrane and hydrolyzed by intracellular esterases to form nonfluorescent 2′,7′-dichlorofluorescein (DCFH), which is rapidly converted to highly fluorescent 2′,7′-dichlorofluorescein (DCF) in the presence of reactive oxygen species. H_2_O_2_ exposure caused a substantial increase in the cellular ROS level indicated in the enhanced the relative green fluorescence (λ_em/DCFDA-H2_=520 nm) intensity of DCFDA-H2 when measured with fluorescence microscopy (Figure 2c & 2d) or flow cytometry (Figure 2e). However, pretreatment with 30 μg mL^-1^ C-Mn_3_O_4_ NPs significantly lowered intracellular ROS level which was reflected in decreased fluorescence (∼ 50% reduction; p<0.001 compared to H_2_O_2_ treated cells) of the probe. The morphological observations in differential interference contrast (DIC) microscopy (Figure 2d) support the results of cell viability and oxidative damage evaluation. The cells pretreated with C-Mn_3_O_4_ NPs prevented the shrinkage and congregation of the cell body due to H_2_O_2_ overexposure and maintained normal cellular architecture.

**Figure 2.**
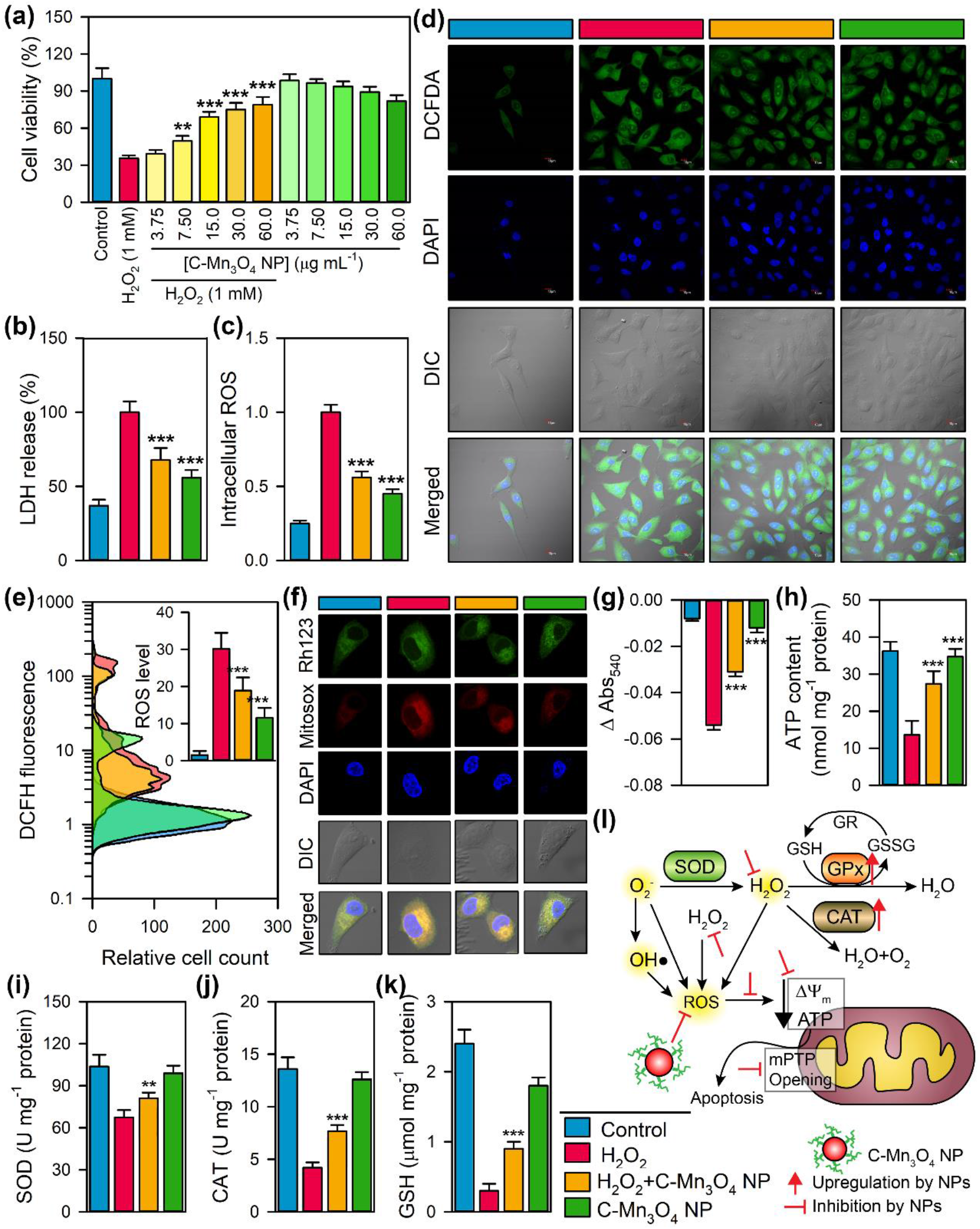
Ability of C-Mn_3_O_4_ NPs in regulation of cellular redox and protection of mitochondria from oxidative damage. (a) Cell viability as measured using MTT. (b) LDH release. (c) Quantification of intracellular ROS estimated from fluorescence microscopy. (d) Fluorescence micrographs of differently stained HEK 293T cells. (e) Intracellular ROS content as measured using flow cytometry using DCFDA. (f) Changes in mitochondrial membrane potential (stained with rhodamie 123) and mitochondrial ROS (stained with Mitosox™) measured using fluorescence imaging. (g) Change in Ca^2+^-induced mPTP opening. (h) ATP content. (i) Superoxide dismutase (SOD) activity. (j) Catalase activity. (k) Reduced glutathione (GSH) content. (l) Schematic representation of the redox homeostasis by C-Mn_3_O_4_ NPs against H_2_O_2_ distress through mitochondrial protection. For all experiments, [H_2_O_2_] =1 mM; [C-Mn_3_O_4_ NPs] =30 μg mL^-1. *, **, ***^ Values differ significantly from H_2_O_2_ treated cells (without treatment) (^***^p < 0.001; ^**^p < 0.01; ^*^p < 0.05).

### C-Mn_3_O_4_ NPs prevent mitochondria, the master redox regulator from H_2_O_2_-induced oxidative damage

Mitochondria despite being the primary source of intracellular ROS and its control is the most susceptible organelle to oxidative damage leading to redox imbalance and cell death [28, 29]. So, to get further insight into the free radical scavenging activity of C-Mn_3_O_4_ NPs we evaluated their protective effect towards mitochondria. Treatment with H_2_O_2_ drastically decreased the ΔΨ_m_, as measured by enhanced rhodamine 123 (Rh123) fluorescence (Figure 2f; Supplementary Figure S2a) along with a burst in mitochondrial ROS production, as indicated by the increased fluorescence of Mito-sox red (Figure 2f; Supplementary Figure S2b). Pretreatment with 30 μg mL^-1^ C-Mn_3_O_4_ NPs significantly restored the ΔΨ_m_ and reduced the mitochondrial ROS. ΔΨ_m_ has a causal relationship with mPTP. The results of Ca^2+^ induced mitochondrial swelling assay indicated that the NPs were effective in preventing the H_2_O_2_ induced mPTP opening (Figure 2g) and maintaining mitochondrial integrity. The mitochondrial membrane depolarization and subsequent opening of mPTP led to a significant fall in the cellular ATP content (Figure 2h). In C-Mn_3_O_4_ NPs pretreated cells, such loss in ATP content was not observed. The opening of mPTP, fall in ΔΨ_m_ and ATP content cumulatively functions as a proapoptotic signal to initiate the cell death pathways. Superoxide dismutase (SOD), catalase (CAT), and glutathione peroxidase (GPx) constitute the intracellular antioxidant defense system that works in consort with mitochondria [30-32]. The accumulation of highly reactive oxygen radicals causes damage to biomolecules in cells and alters enzyme activities [33-35]. Hence, we extended our study towards evaluating the effect of H_2_O_2_ and C-Mn_3_O_4_ NPs in the ROS regulatory network. H_2_O_2_ exposure significantly reduced the activity of SOD, CAT, and GPx resulting in a decrease of the reducing pool of cellular thiol constituents (e.g., GSH) (Figure 2i-2k). Pretreatment with C-Mn_3_O_4_ NPs significantly attenuated the damage. In cells treated with C-Mn_3_O_4_ NPs alone, none of the detrimental effects were observed.

Thus, our cellular studies indicate that C-Mn_3_O_4_ NPs possess the distinctive property of scavenging intracellular ROS, inhibiting apoptotic trigger, preventing loss of antioxidant enzymes and maintaining high cell viability by acting as a protector of mitochondria, the master regulator of cellular redox equilibrium (Figure 2l schematically summarizes the whole sequence).

### C-Mn_3_O_4_ NPs attenuate glomerular and tubulointerstitial damage in CKD mice

There is always a gap in the efficacies of a pharmacological agent tested between cellular and animal model. The limited bioavailability, nonspecific distribution, or unwarranted metabolism often restricts the use of a cytoprotective agent *in vivo* [36-38]. Thus, we evaluated the potential of C-Mn_3_O_4_ NPs in the treatment of cisplatin-induced C57BL/6j mice, a well-known animal model for testing therapeutic interventions against CKD [39-41]. Chronic administration of cisplatin resulted in significant mortality (∼40% compared to control) (Figure 3a). The fourfold higher blood urea nitrogen (BUN) content (Figure 3b), threefold higher GFR (Table 1), fourfold higher urinary albumin excretion (albuminuria) (Figure 3c) and high urine albumin to creatinine ratio (ACR) (Table 1) along with significantly increased serum urea (Figure 3d) and creatinine (Figure 3e) illustrated induction of proteinuria and notable damage to the renal function of mice, the two hallmarks of CKD [42-44]. Treatment with C-Mn_3_O_4_ NPs (0.25 mg kg^-1^ body weight (BW)) considerably reduced BUN, GFR, urinary albumin, ACR, serum urea, and creatinine (Figure 3b-3e; Table 1). The treatment also improved survivability (Log-rank *p*=0.022; hazards ratio 2.62) (Figure 3a). Treatment with citrate (the functionalization group) was unable to reduce any of the aforementioned parameters (Supplementary Figure S3), confirming the observed effects solely due to the conjugated nanomaterial. Cisplatin intoxication caused weight loss in mice (Figure 3f), suggestive of the systemic toxicity that frequently arises in individuals receiving this anticancer drug. Animals treated with C-Mn_3_O_4_ NPs were capable of mitigating weight loss. Next, we examined the external morphology of isolated kidneys from each group. The kidneys from the cisplatin exposed group were deviated from the usual darkish brown to a pale brown color with a rough and uneven surface (Figure 3g). The kidney to body weight ratio (i.e, kidney index) was also significantly higher (i.e., edema) in cisplatin-treated animals (2.1±0.2 compared to 1.5±0.1 mg g^-1^ of control, p<0.001; Figure 3h). Subsequent treatment with C-Mn_3_O_4_ NPs overturned the observed changes in morphology and kidney index.

**Table 1.**
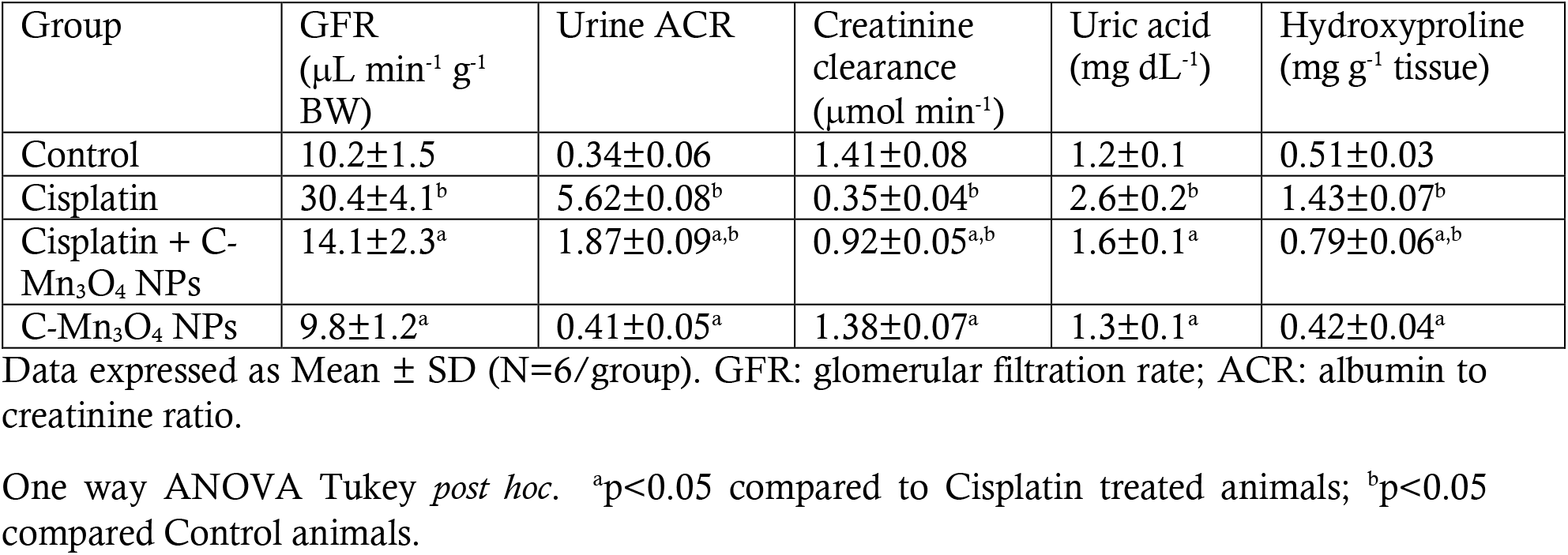
Effect of C-Mn_3_O_4_ NPs on nephrotoxic biomarkers.

| Group | GFR<br>($\mu\text{L min}^{-1} \text{g}^{-1}$<br>BW) | Urine ACR | Creatinine<br>clearance<br>($\mu\text{mol min}^{-1}$) | Uric acid<br>(mg dL<sup>-1</sup>) | Hydroxyproline<br>(mg g<sup>-1</sup> tissue) |
| --- | --- | --- | --- | --- | --- |
| Control | 10.2±1.5 | 0.34±0.06 | 1.41±0.08 | 1.2±0.1 | 0.51±0.03 |
| Cisplatin | 30.4±4.1<sup>b</sup> | 5.62±0.08<sup>b</sup> | 0.35±0.04<sup>b</sup> | 2.6±0.2<sup>b</sup> | 1.43±0.07<sup>b</sup> |
| Cisplatin + C-Mn<sub>3</sub>O<sub>4</sub> NPs | 14.1±2.3<sup>a</sup> | 1.87±0.09<sup>a,b</sup> | 0.92±0.05<sup>a,b</sup> | 1.6±0.1<sup>a</sup> | 0.79±0.06<sup>a,b</sup> |
| C-Mn<sub>3</sub>O<sub>4</sub> NPs | 9.8±1.2<sup>a</sup> | 0.41±0.05<sup>a</sup> | 1.38±0.07<sup>a</sup> | 1.3±0.1<sup>a</sup> | 0.42±0.04<sup>a</sup> |

Data expressed as Mean ± SD (N=6/group). GFR: glomerular filtration rate; ACR: albumin to creatinine ratio.
One way ANOVA Tukey *post hoc*. <sup>a</sup>p<0.05 compared to Cisplatin treated animals; <sup>b</sup>p<0.05 compared Control animals.

**Figure 3.**
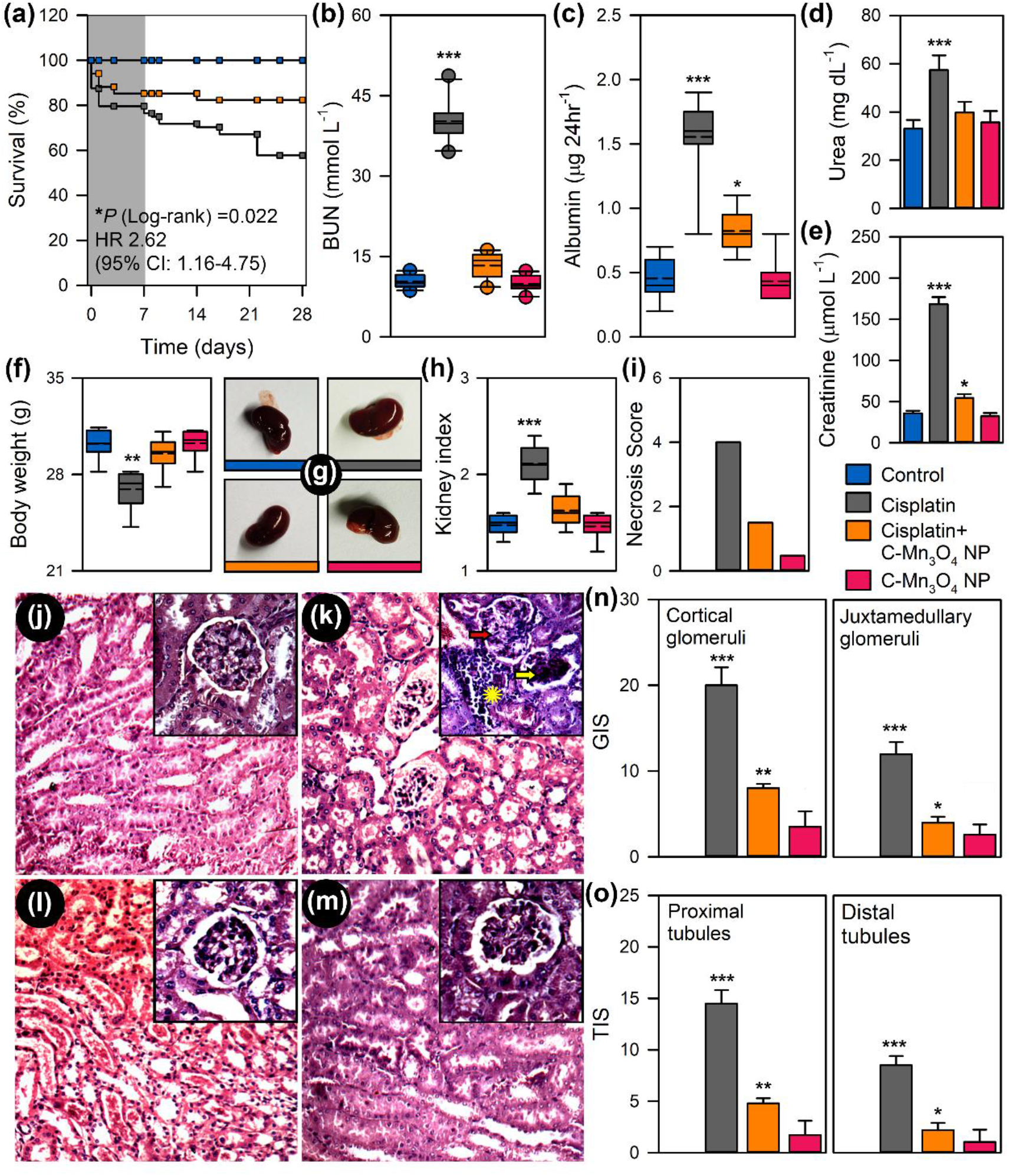
Efficacy of C-Mn_3_O_4_ NPs in reversal of CKD in animal model. (a) Kaplan-Meir survival analysis curve. The darker shaded area represents co-treatment period. (b) Blood urea nitrogen (BUN) content. (c) Urinary albumin excretion as an indicator of albuminuria, hallmark of CKD. (d) Serum urea concentration. (e) Serum creatinine level. (f) Body weight at the end of experimental period. (g) Photographs of kidneys incised after experimental period. (h) Kidney index, defined as kidney to body weight ratio (mg g^-1^). (i) Necrosis score as per the observation of expert clinical pathologist. (j-m) Hematoxylin and eosin stained liver sections. j: Control; k: Cisplatin; l: Cisplatin+C-Mn_3_O_4_ NPs; m: C-Mn_3_O_4_ NPs. Inset shows magnified image of single glomerulus. Red arrow: segmental glomerulosclerosis; Yellow arrow: global glomerulosclerosis; Yellow star: mononuclear infiltration (n) Glomerular injury score (GIS). (o) Tubular injury score (TIS). Data are expressed as Mean ± SD. *N*=6. ^*, **, ***^ Values differ significantly from control group (without treatment) (^***^p < 0.001; ^**^p < 0.01; ^*^p < 0.05).

Hematoxylin and eosin-stained kidney sections of the control and C-Mn_3_O_4_ NP treated groups showed normal histologic features (Figure 3j & 3m) with negligible necrosis score (Figure 3i). The kidney sections from cisplatin intoxicated mice displayed several pathological features of CKD like focal segmental as well as global glomerulosclerosis along with interstitial fibrosis, diffused thickening of the capillary walls, glomerular hyalinosis, dilated or collapsed Bowman’s space and glomerular retraction (Figure 3k). Tubular atrophy, dilation of cortical tubules, increased mesangial matrix, obliteration of capillaries, necrosis, vacuolization, and interstitial mononuclear infiltration were the other features observed in this group. Treatment with C-Mn_3_O_4_ NPs notably reduced focal glomerular necrosis (Figure 3l). However, sparse tubular changes like vacuolization, dilation, mild mononuclear infiltration, and detachment of epithelial cells were observed in this group. Overall, C-Mn_3_O_4_ NPs were able to efficiently revert the marked detrimental changes in the renal architecture of CKD animals. The histological observations are quantitatively reflected in the necrosis score (Figure 3i), glomerular injury score (GIS; Figure 3n), and tubular injury score (TIS; Figure 3o).

Previous studies and our histological observations suggested an association between renal fibrosis and CKD [45-47]. So, we measured the renal hydroxyproline content, a byproduct of collagen metabolism and biochemical marker of fibrosis. The results indicate almost a threefold increase in the hydroxyproline content (1.43±0.07 compared to 0.51±0.03 mg gm^-1^ tissue of control; p<0.001) in the cisplatin intoxicated group (Table 1). In accordance with the histological findings, treatment with C-Mn_3_O_4_ NPs markedly reduced the hydroxyproline content (0.79±0.06 mg gm^-1^ tissue; p<0.001 compared to cisplatin treated ones), suggesting a decrease in fibrotic damage (Table 1).

### C-Mn_3_O_4_ NPs augment the intracellular antioxidant defense system

Oxidative stress proved to be one of the major causes of cisplatin-induced nephrotoxicity [48-50]. Therefore, we tried to ascertain whether C-Mn_3_O_4_ NPs contributed to nephroprotection by ameliorating oxidative stress. Signs of ROS mediated damage including lipid peroxidation (in terms of thiobarbituric acid reactive substances, TBARS), reduction in cellular GSH pool, and inhibition of antioxidant enzyme activity were estimated. Exposure to cisplatin markedly increased the level of TBARS (Figure 4a) and oxidative glutathione along with a reduction in GSH concentration. Furthermore, it inhibited the antioxidant actions of enzymes like SOD, CAT, and GPx (Figure 4b-4d). In consensus with our cellular studies, C-Mn_3_O_4_ NPs rescued the detrimental pleiotropic effects of increased ROS while maintaining the normal signaling circuitry.

**Figure 4.**
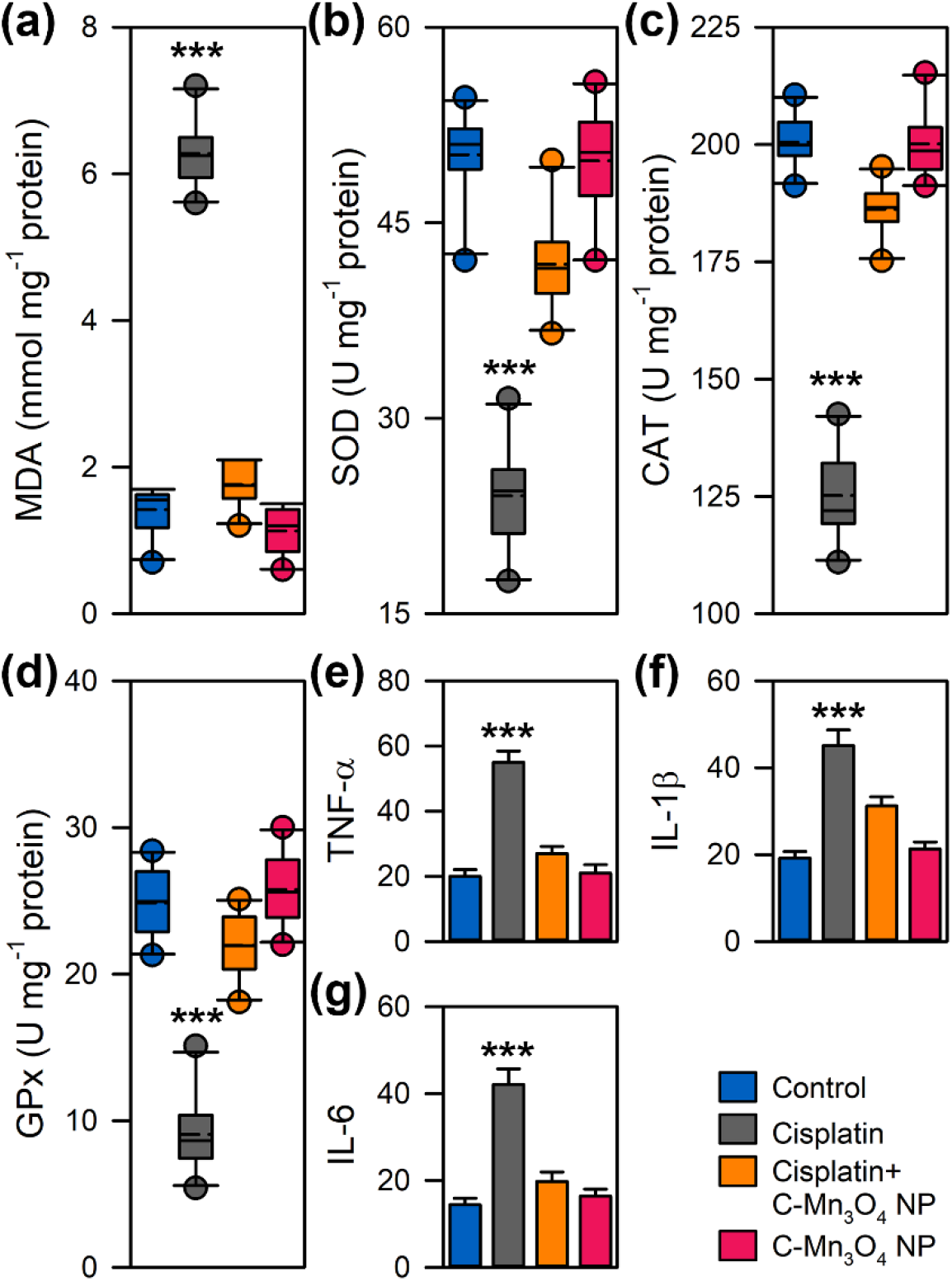
Effect of C-Mn_3_O_4_ NPs in protection of intracellular redox regulatory network and inhibition of anti-inflammatory response. (a) Extent of lipid peroxidation (MDA, malonaldehyde content) measured in terms of thiobarbituric acid reactive substances (TBARS). (b) Superoxide dismutase (SOD) activity. (c) Catalase activity. (d) Gluthione peroxidase (GPx) activity. (e) Tumor necrosis factor-α level. (f) Interleukin-1β level. (g) Interleukin-6 level. MDA, SOD, CAT and GPx were estimated from kidney homogenate. TNF-β, IL-1β and IL-6 were measured from serum. Data are expressed as Mean ± SD. *N*=6. ^*, **, ***^ Values differ significantly from control group (without treatment) (^***^p < 0.001; ^**^p < 0.01; ^*^p < 0.05).

### C-Mn_3_O_4_ NPs reduce renal inflammation

Macrophage infiltration in the kidney and subsequent rise in the plasma concentrations of pro-inflammatory cytokines like TNF-α are well-known features of CKD [51-53]. We found significant increases in plasma concentrations of TNF-α, IL-1β, and IL-6 in cisplatin-induced animals (Figure 4e-4g). Treatment with C-Mn_3_O_4_ NPs resulted in a notable decrease in the cytokine levels. No difference was observed between the C-Mn_3_O_4_ NP treated and the control groups.

### C-Mn_3_O_4_ NPs alleviate mitochondrial damage in CKD mice

Considering the inevitable role of mitochondria in the pathogenesis of CKD [26, 54-59] and results of our *in cellulo* observations that C-Mn_3_O_4_ NPs protects mitochondria from H_2_O_2_ induced oxidative damage, we assessed the role of mitoprotection in the therapeutic efficacy of C-Mn_3_O_4_ NPs in animals. Ca^2+^-induced renal mPTP opening is one of the salient features of CKD [26, 60]. Our data clearly show that the mitochondria isolated from the cisplatin intoxicated group were more sensitive towards Ca^2+^ manifested into a sharp decrease in 540 nm absorbance (Figure 5a). Treatment with C-Mn_3_O_4_ NPs inhibited mPTP opening and maintained membrane integrity. ΔΨ_m_ and ATP content declined significantly as a result of cisplatin administration (Figure 5b & 5c). These were accompanied by an increase in cytochrome c oxidase activity (Figure 5d) and a reduction in dehydrogenase activity (Figure 5e). The alterations were upended by C-Mn_3_O_4_ NP treatment. Thus cisplatin-induced renal damage triggered the opening of mPTP, the decline in ΔΨ_m_, and induction of mitochondrial swelling that resulted in the release of cytochrome c in the cytosol leading to apoptosis. The ladder-like DNA fragmentation, a hallmark of apoptosis, was evident in the case of cisplatin-treated diseased groups (Figure 5f). Whereas, treatment with C-Mn_3_O_4_ NPs notably protected the mitochondria, inhibited cell death and decreased the extent of DNA fragmentation. Figure 5g schematically represents the entire phenomena of redox-mediated nephroprotection by C-Mn_3_O_4_ NPs.

**Figure 5.**
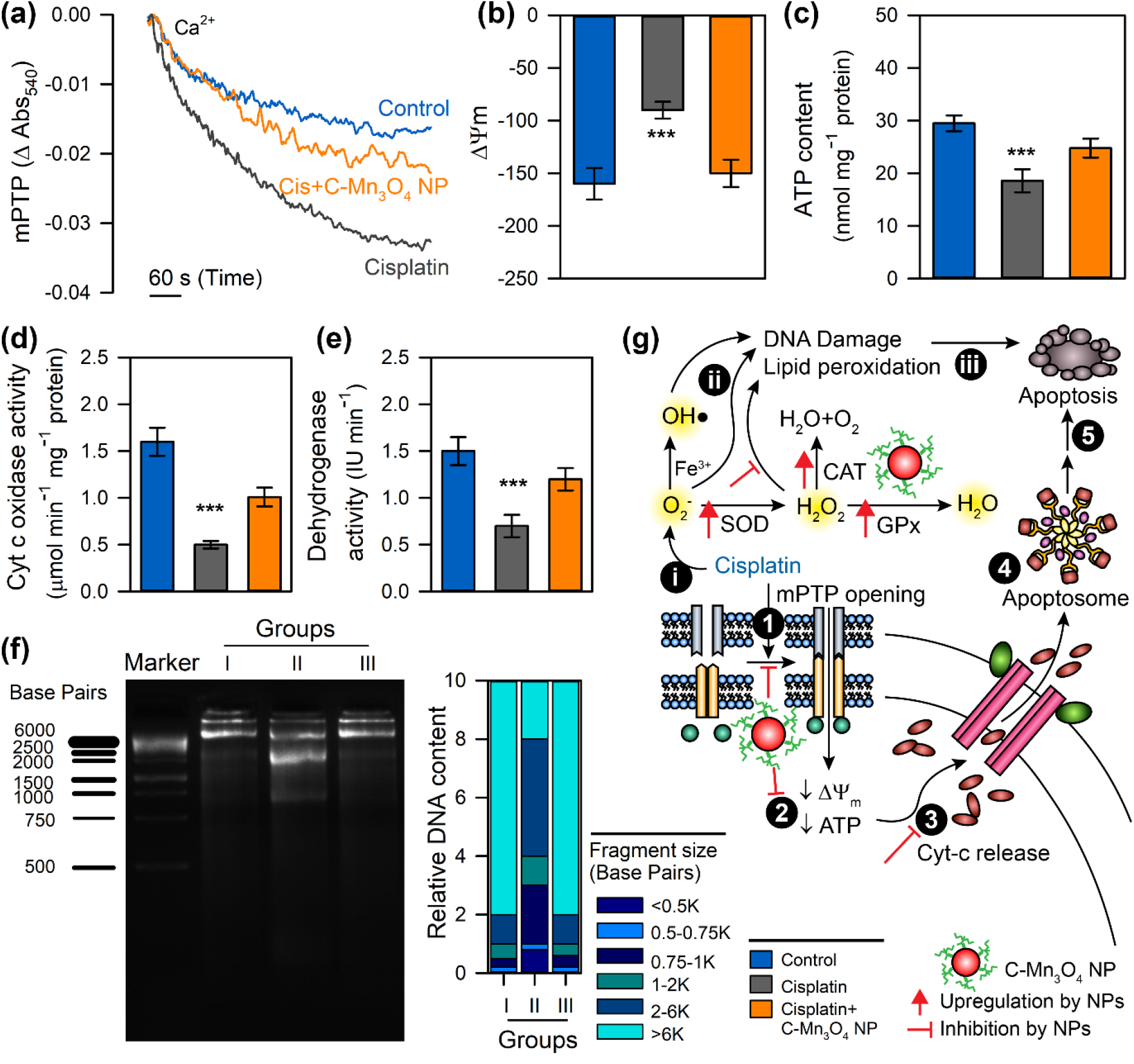
Efficacy of C-Mn_3_O_4_ NPs in protection of mitochondria, the master redox regulator in mice. (a) Ca^2+^ induced mPTP opening. (b) Mitochondrial membrane potential measured using JC-1. (c) ATP content. (d) Cytochrome c oxidase (complex IV) activity in isolated mitochondria. (e) Succinate dehydrogenase (SDH, complex II) activity in isolated mitochondria. (f) DNA fragmentation level as a result of oxidative damage measured using agarose gel electrophoresis. Group I: Control; Group II: Cisplatin; Group II: Cisplatin+C-Mn_3_O_4_ NPs. (g) Schematic overview of the comprehensive mechanism of action of C-Mn_3_O_4_ NPs as a redox medicine against cisplatin induced CKD. The numbers in the black circles indicate the sequence of events. Data are expressed as Mean ± SD. *N*=6. ^*, **, ***^ Values differ significantly from control group (without treatment) (^***^p < 0.001; ^**^p < 0.01; ^*^p < 0.05).

## DISCUSSION

In this study, we determined whether C-Mn_3_O_4_ NPs could be used as a redox medicine to treat CKD, an important clinical question considering the high prevalence of the disease and the non-availability of effective medication. CKD is defined as the progressive and irreversible loss of renal function characterized by reduced glomerular filtration rate (GFR), increased urinary albumin excretion (albuminuria), or both [42, 44, 61]. Our results present evidence that treatment with C-Mn_3_O_4_ NPs significantly improved renal function, glomerular and tubulointerstitial injury, cellular antioxidant defense network in line with inhibition of pro-inflammatory immune response and attenuation of mitochondrial dysfunction in response to the cisplatin toxicity. Our cellular and animal studies further enlightened the role of unique mitoprotective as well as redox modulatory activity of C-Mn_3_O_4_ NPs in the therapeutic mechanism.

Mitochondria have long been recognized for their canonical roles in cellular respiration and energy production [26]. Recently, they have emerged as the master regulator of a spectrum of molecular pathways including biosynthesis of macromolecules, maintenance of cellular redox equilibrium, calcium homeostasis, inflammation, and cell death [62-66]. Thus, mitochondria are poised to play a pivotal role in the functioning of the kidney, an organ with high energy demand, and rich in mitochondria, second only to the heart [56]. Our findings that C-Mn_3_O_4_ NPs maintain cellular redox homeostasis through the prevention of mPTP opening and ATP depletion discloses a key redox-mediated nephroprotective mechanism. Virtually, the renal proximal tubules are exclusively dependent on ATP generated by mitochondrial oxidative phosphorylation and are therefore vulnerable to the oxidative distress due to mitochondrial damage [48, 67]. Cisplatin accrues in mitochondria and reduces the activity of all four respiratory complexes (I–V) involved in the electron transport chain, thereby a surge in mitochondrial ROS formation takes place along with mPTP opening, membrane depolarization and impairment in ATP production, leading to cell death [68, 69]. The mitotoxic mechanism of cisplatin essentially mimics the pathogenesis of CKD, thus an efficient reversal of damage in this rodent model is supposed to reflect the possible effects of a compound in higher animals. Data from our cellular as well as animal studies provide sufficient evidence that C-Mn_3_O_4_ NPs prevent mitochondrial ROS surge, prevent loss of membrane potential, inhibit mPTP opening, and stops ATP depletion, thereby prevents mitochondrial dysfunction, cellular redox imbalance, and tubular or glomerular cell death. As a result, the markers of CKD like increased BUN, plasma creatinine, serum urea, and glomerular filtration rate (GFR) returns to homeostatic condition.

This study provides a piece of direct evidence that C-Mn_3_O_4_ NPs can scavenge ROS, particularly H_2_O_2_ the longest living one in the cellular *milieu*. It also proves the ability of the NPs in the prevention of mPTP from opening and subsequently maintenance of mitochondrial structure and function. However, it is not clear whether ROS scavenging protects mitochondria or mitochondrial integrity which results in ROS depletion. Several studies showed that a compound with the sole property of free radical scavenging cannot be effective in the reversal of oxidative stress-related diseases in higher animals because the antioxidant property is not sustainable and the compound becomes inactive after one reaction [70-72]. So, to mitigate the burst of ROS, the intracellular concentration required for a conventional antioxidant molecule is enormous and practically hard to achieve [70-72]. Thus, to become an effective medicine a compound should have some additional mechanisms that will ensure the sustainable effect in the therapeutic regime. Therefore, considering the causal relationship between mitochondria and cellular redox homeostasis and the efficacy of C-Mn_3_O_4_ NPs in the treatment of multifaceted diseases like CKD, we propose that both the mechanisms (i.e., ROS scavenging and mitochondrial protection) simultaneously take place.

The findings that C-Mn_3_O_4_ NPs can accelerate the revival of proximal tubule epithelium embodies a crucial nephroprotective function mediated by the nanoparticles. Kidneys show higher regenerative property following tubulointerstitial damage [48]. The proliferation of a subset of sublethally damaged, yet surviving, proximal tubule cells contribute to the regenerative property of the kidney [73]. The acceleration of this process is sufficient to confer nephroprotection [74, 75]. As revealed in our histological findings, the recovery rate of these cells in C-Mn_3_O_4_ NP treated cisplatin exposed animals is significantly faster and efficient than the auto-recovery. Several mechanisms can be proposed for the enhanced proliferation by C-Mn_3_O_4_ NPs. The restoration of structural and functional integrity of mitochondria and recovery of respiratory complexes may contribute towards the increased proliferation. It is well known that the mitochondrial electron transport chain (ETC) has a crucial role in cell proliferation through regulation of ATP generation, and supply of energy to proliferative pathways [76, 77]. Previous studies have demonstrated that mutations in ETC genes, or presence of ETC complex inhibitors causes a reduction in ATP synthesis obstructing progression through cell cycle leading to a blockage in proliferation [78-80]. Thus, the mitoprotective activity of C-Mn_3_O_4_ NPs may have played a significant role in the revival of tubulointerstitial epithelial cells, in turn protecting the renal architecture. Additionally, ROS scavenging by C-Mn_3_O_4_ NPs may boost the proliferation because oxidative distress in proximal tubules causes cell cycle arrest and impedes cell-cycle progression [27].

The role of intracellular redox regulation through mitoprotection in the therapeutic action of C-Mn_3_O_4_ NPs opens up further avenues for the treatment of several unmet diseases like diabetic nephropathy, neurodegeneration (e.g., Parkinson’s, Huntington’s, Alzheimer’s, multiple sclerosis), cardiovascular disorders, obesity, etc. where pathogenesis is very much dependent upon mitochondrial damage and associated redox imbalance [81-85]. Although in our study C-Mn_3_O_4_ NPs did not show any adverse side effect, a detailed study on toxicity, bio-distribution, and pharmacokinetics will greatly enhance the knowledge about its *in vivo* behavior. Furthermore, a detailed molecular study analyzing the genome and metabolome of C-Mn_3_O_4_ NP treated animals may enlighten its ability to interfere in other pathogenesis pathways. As a result, the nano-drug could be repurposed for other therapies too.

## CONCLUSION

There are very few published articles that utilize the promising redox regulatory approach for treatment of chronic diseases like CKD. On the other hand, several chronic kidney diseases are reported to be due to redox imbalance in mitochondria. Our study suggests that C-Mn_3_O_4_ NPs could be an efficient redox medicine to attenuate renal injury and tubuleintestinal fibrosis as evidenced by the improved renal functions, reduction in biochemical markers of nephrotoxicity, reduced fibrotic content, and downregulated proinflammatory cytokines. The molecular mechanism involves regulation of the redox balance through synchronization of the causal relationship between mitoprotection and ROS scavenging by C-Mn_3_O_4_ NPs. The findings highly suggested the translational potential of C-Mn_3_O_4_ NPs as a redox nanomedicine for treating CKD in the clinic.

## MATERIALS AND METHODS

### Synthesis of C-Mn_3_O_4_ NPs

A template or surfactant-free sol-gel method was followed for the synthesis of bulk Mn_3_O_4_ NPs at room temperature and pressure [21]. To functionalize the nanoparticles with ligand citrate, the as-prepared Mn_3_O_4_ NPs (∼20 mg mL^-1^) were mixed extensively with citrate (Sigma, USA) solution of (pH 7.0, 0.5 M) for 15 h in a cyclomixer. The time for mixing was carefully adjusted to have a nanomaterial in the size range of 4-6 nm. Non-functionalized larger NPs were removed using a syringe filter (0.22 μm).

### Characterization techniques

TEM and HRTEM images were acquired using an FEI TecnaiTF-20 field emission HRTEM operating at 200 kV. Sample preparation was done by drop-casting of C-Mn_3_O_4_ NP solution on 300-mesh amorphous carbon-coated copper grids (Sigma, USA) and allowed to dry overnight at room temperature. XRD patterns were obtained by employing a scanning rate of 0.02 s^-1^ in the 2*θ* range from 10 to 80 by a PANalytical XPERT PRO diffractometer equipped with Cu Ka radiation (at 40 mA and 40 kV). FTIR (JASCO FTIR-6300, Japan) was used to confirm the covalent attachment of the citrate molecules with the Mn_3_O_4_ NPs. For FTIR studies, powdered samples were blended with KBr powder and pelletized. KBr pellets were used as a reference to make the background correction.

### H_2_O_2_ scavenging by C-Mn_3_O_4_ NPs

The ability of C-Mn_3_O_4_ NPs to prevent H_2_O_2_ mediated degradation of sodium containing dye RB (Sigma, USA) was used as an indicator of its H_2_O_2_ scavenging activity. Addition of H_2_O_2_ (10 mM) in the aqueous solution of RB (3.5 μM) leads to decolourization of the dye reflected into a decrease in absorbance (λ_max_=550 nm). Presence of C-Mn_3_O_4_ NPs (50 μg mL^-1^) in the reaction mixture reduced degradation. All absorbance measurements were performed using Shimadzu UV-Vis 2600 spectrometer (Tokyo, Japan).

### Culture of Human Embryonic Kidney Cells (HEK 293)

HEK 293 cells were maintained at 37°C in 5% CO_2_ in RPMI 1640 growth medium (Himedia, India) that contained 10% foetal bovine serum (Invitrogen, USA), L-glutamine (2 mM), penicillin (100 units mL^-1^), and streptomycin (100 ng mL^-1^) (Sigma, USA). Before experimentation, the cells were washed twice and incubated with RPMI 1640 medium (FBS, 0.5%) for 1 h and then treated as described in the figure legends.

### Measurement of cell viability

Cell viability was assessed by MTT and LDH assay. All cell lines were plated in 96-well plates at a density of 1×10^3^ cells/well and cultured overnight at 37°C. The treatments were performed as described in the figure legends. Next, MTT (5 mg mL^-1^; Himedia, India) was added to each well, with a final concentration of ∼0.5 mg mL^-1^, and the cells were cultured for 4 h at 37°C in a 5% CO_2_ atmosphere. Resultant purple formazan was dissolved by the addition of 10% sodium dodecyl sulfate (Sigma, USA) and the absorbance was read at 570 nm and 630 nm using a microplate reader (BioTek, USA). LDH release was analyzed using a colorimetric LDH cytotoxicity assay (Himedia, India) following the manufacturer’s instructions. Three independent experiments were performed in each case.

### Measurement of intracellular ROS

The formation of intracellular ROS was measured with the DCFH-DA method using both FACS and confocal microscopy. For FACS, after treatment cells were trypsinized, washed with 1X PBS and stained with DCFDA-H2 (15 μM; Sigma, USA) for 10 mins at 30°C in dark. Ten thousand events were analyzed by flow cytometry (FACS Verse, Beckton Dickinson, SanJose, USA) and the respective mean fluorescence intensity (in FL1 channel, set with a 530/30 nm bandpass filter) values were correlated with the ROS levels. For confocal microscopy, 5000 cells were seeded and treated with agents described earlier. Post-treatment cells were stained with DCFDA-H2 (5 μM) at 37°C in dark and images were acquired with a confocal microscope (Olympus IX84, Japan). All parameters (pinhole, contrast, gain, and offset) were held constant for all sections in the same experiment.

### Mitochondrial superoxide and ΔΨ_m_ detection

After treatment with the drug or vehicle, mitochondrial superoxide production was visualized using MitoSOX Red, a mitochondrial superoxide indicator (Thermofisher, USA). The ΔΨ_m_ in intact cells was assessed by confocal microscopy using Rh 123. Cells were loaded with MitoSOX Red (0.5 μM) or Rh123 (1.5 μM) for 10 min at 37°C and imaged using a confocal microscope (Olympus IX84, Japan). All parameters (pinhole, contrast, gain, and offset) were held constant for all sections in the same experiment. ImageJ (http://imagej.nih.gov/ij/) was used to quantify area normalized florescence intensities from the confocal images.

### Animals and treatment

Healthy non-diabetic C57/6j mice of both sexes (8–10 weeks old, weighing 27±2.3 g) were used in this work. Animals were maintained in standard, clean polypropylene cages (temperature 21 ± 1°C; relative humidity 40–55%; 1:1 light and dark cycle). Water and standard laboratory pellet diet for mice (Saha Enterprise, Kolkata, India) were available *ad libitum* throughout the experimental period. All mice were allowed to acclimatize for 2 weeks before the treatment. The guideline of Committee for the Purpose of Control and Supervision of Experiments on Animals, New Delhi, India, was followed and the study was approved by the Institutional Animal Ethics Committee (Ethical Clearance No. - 05/S/UC-IAEC/01/2019).

The mice were randomly divided into five groups (n = 16 for each group): (1) control; (2) cisplatin; (3) cisplatin + C-Mn_3_O_4_ NPs; and (4) C-Mn_3_O_4_ NPs; (5) cisplatin + citrate. The experimental model of CKD was established according to the previous description. In brief, for induction of CKD, we used 8 mg kg^-1^ BW cisplatin (i.p.) in each alternative day for 28 days. After induction, we treated C-Mn_3_O_4_ NPs at 0.25 mg kg^-1^ BW (i.p.) for another 28 days. There was an overlap of 7 days between induction and treatment. Citrate was used at a dose of 0.25 mg kg^-1^ BW for 28 days. All doses were finalized based on reported literature and pilot experimentation. As citrate treatment did not improve the kidney function, data are not represented in the main manuscript.

### Biochemical evaluations

Whole blood samples from treated mice were collected from retro-orbital sinus plexus and centrifuged at 2000 ×g for 20 min to separate the serum. Urine samples were collected in metabolic cages during 24-h fasting conditions. Biochemical evaluations were performed using commercially available kits (Autospan Liquid Gold, Span Diagnostic Ltd., India) following the protocol described by respective manufacturers. GFR was estimated by the determination of urinary excretion of fluorescein-labeled inulin (FITC–inulin).

### Histological examination

After incision, kidney tissues from mice were fixed with 4% paraformaldehyde, embedded in paraffin, and cut into 5 μm thick section. After de-waxing and gradual hydration with ethanol (Merck, USA), kidney sections were stained with hematoxylin and eosin (SRL, India). The sections were then observed under an optical microscope (Olympus, Tokyo, Japan).

### Renal hydroxyproline measurement

For the measurement of renal hydroxyproline content, a previously described method was used [31]. In brief, snap-frozen kidney specimens (200 mg) were weighed, hydrolyzed in HCl (6M; Merck, USA) for 12 h at 100°C. Next, they were oxidized with Chloramine-T (SRL, India). Next, Ehrlich reagent (Sigma, USA) was added which resulted in the formation of a chromophore. Absorbance was measured at 550 nm. Data were normalized to kidney wet weight.

### Renal homogenate preparation

Samples of kidney tissue were collected, homogenized in cold phosphate buffer (0.1 M; pH 7.4), and centrifuged at 10,000 r.p.m. at 4°C for 15 min. The supernatants were collected for further experimentation.

### Assessment of lipid peroxidation & hepatic antioxidant status

The supernatants obtained in the previous stage were used to measure the activity of SOD, CAT, GPx, and GSH as well as the content of lipid peroxidation (MDA). Lipid peroxidation was determined in TBARS formation using a reported procedure [30]. SOD (Sigma, MO, USA), CAT (Abcam, Germany), and GPx activities (Sigma, MO, USA) were estimated using commercially available test kits following protocols recommended by respective manufacturers. The renal GSH level was determined by the method of Ellman with trivial modifications [86].

### Mitochondria isolation and mitochondrial function determination

Mitochondria were isolated from mouse kidneys following the method described by Graham [87] with slight modifications. In brief, kidneys were excised and homogenized in kidney homogenization medium containing 225 mM D-mannitol, 75 mM sucrose, 0.05 mM EDTA, 10 mM KCl, 10 mM HEPES (pH 7.4). The homogenates were centrifuged at 600 ×g for 15 min and the resulting supernatants were centrifuged at 8500 ×g for 10 min. The pellets were washed thrice and resuspended in the same buffer. All procedures were performed at 4°C.

Mitochondrial function was evaluated by determining ΔΨ_m_ using JC-1 (Sigma, MO, USA), ATP production (Abcam, Germany), and the activities of mitochondrial complexes succinate dehydrogenase and cytochrome c oxidase. mPTP opening was measured in terms of mitochondrial swelling by monitoring the decrease in absorbance at 540 nm after the addition of CaCl_2_ (100 mM).

### Agarose gel electrophoresis for DNA fragmentation

Total renal DNA was isolated following a standard procedure [88]. DNA fragmentation was assessed using agarose gel electrophoresis. In a typical procedure, renal DNA (5.0 μg) was loaded on 1.5% agarose gel stained with ethidium bromide. Electrophoresis was carried out for 2 h at 90 V, and the resultant gel was photographed under UV transillumination (InGenius 3 gel documentation system, Syngene, MD, USA).

### Statistical analysis

All numerical data are expressed as mean ± standard deviation (SD) unless otherwise specified. One-way analysis of variance (ANOVA) followed by Tukey’s *post hoc* multiple comparison test was implemented to compare various parameters between the groups using a commercially available software GraphPad Prism (version 5.00, GraphPad Software, CA, USA). p < 0.05 was considered significant.

## ACKNOWLEDGMENTS

MD thanks University Grants Commission (UGC), Govt. of India for Junior Research Fellowship. SKP thanks the Indian National Academy of Engineering (INAE) for the Abdul Kalam Technology Innovation National Fellowship, INAE/121/AKF. The authors thank the DBT (WB)-BOOST scheme for the financial grant, 339/WBBDC/1P-2/2013.

## CONFLICT OF INTEREST

The authors disclose no conflict of interest.

## AUTHOR CONTRIBUTION

AA, MB and SKP designed the experiments. AA, SM, and PB did synthesis, characterization and *in vitro* studies. TC, MB performed cellular studies. AA, SD, and MD conducted animal experiments. AD performed the histological studies and analysis. AA, HA, JTA, AS, SAA, MB and SKP discussed and analyzed the results. AA wrote the manuscript and all authors contributed towards writing of the final version.

## Notes

### Competing Interest Statement

The authors have declared no competing interest.

