## Supplementary Information for "Redox Nanomedicine Cures Chronic Kidney Disease (CKD) by Mitochondrial Reconditioning"

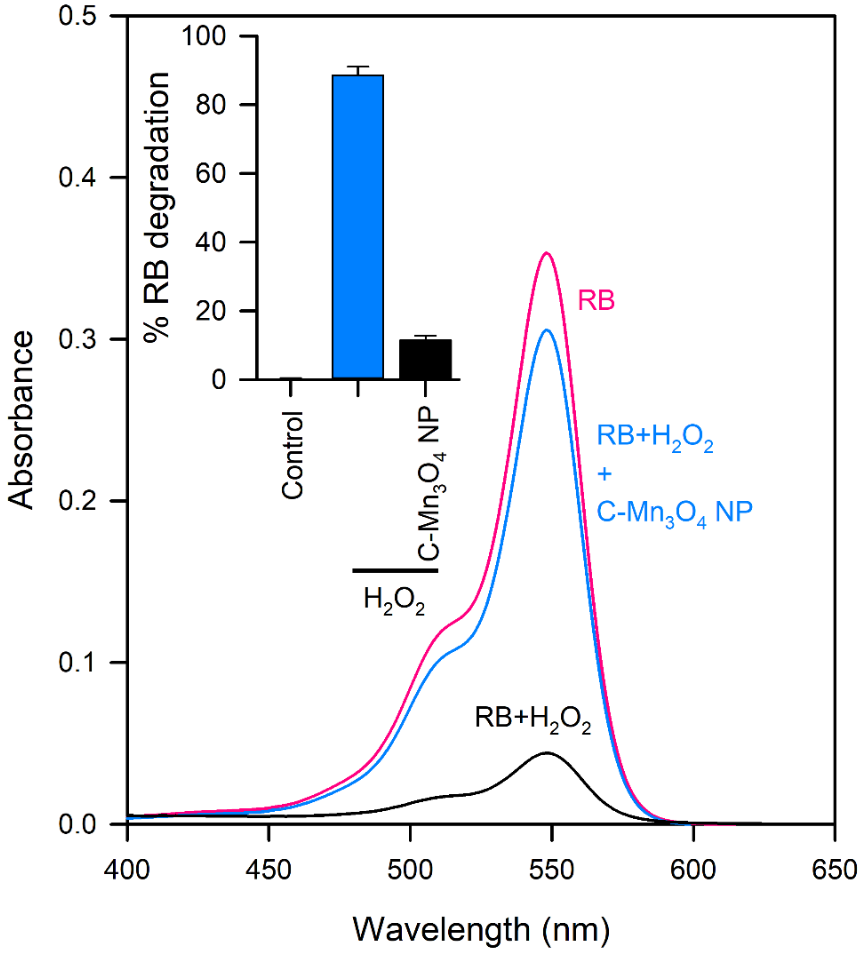


**Supplementary Figure S1. $H_2O_2$ scavenging activity of C-$Mn_3O_4$ NPs using Rose Bengal (RB) assay.** The RB undergoes oxidative degradation upon interaction with $H_2O_2$ as indicated in significantly decreased absorbance spectra. In presence of C-$Mn_3O_4$ NPs, $H_2O_2$ cannot degrade RB due to radical scavenging activity of the NPs. The inset shows the percentage of RB degradation by $H_2O_2$ in absence and presence of C-$Mn_3O_4$ NPs.

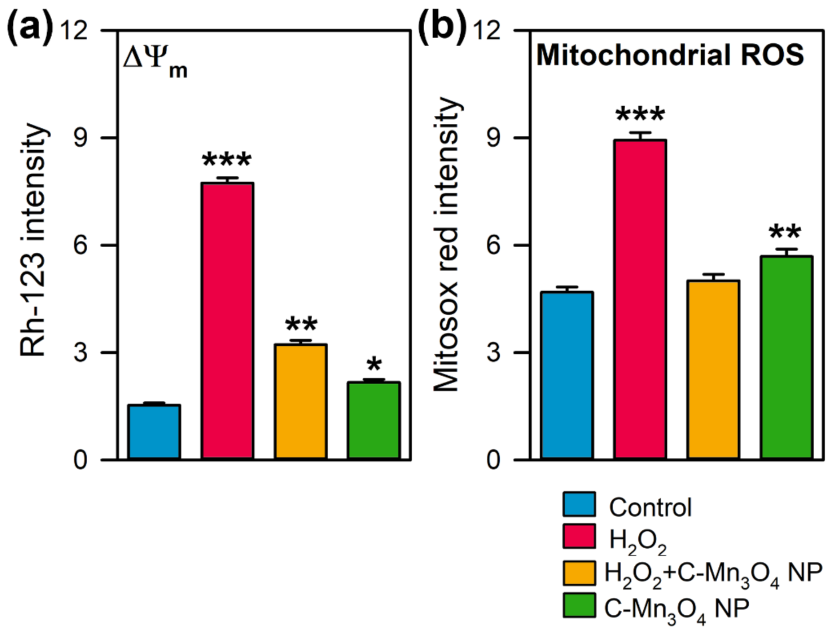


**Supplementary Figure S2. Quantification of fluorescence intensity (Figure 2f) using ImageJ.** (a) Intensity of Rh-123 as a marker of mitochondrial membrane potential ($\Delta\Psi_m$). More intensity indicates increased depolarization. (b) Intensity of Mitosox red™ as a marker of mitochondrial ROS.

All measurements are in arbitrary unit and expressed as Mean ± SD (N=5). *, **, *** Values differ significantly from cells without treatment (***$p < 0.001$; **$p < 0.01$; *$p < 0.05$).

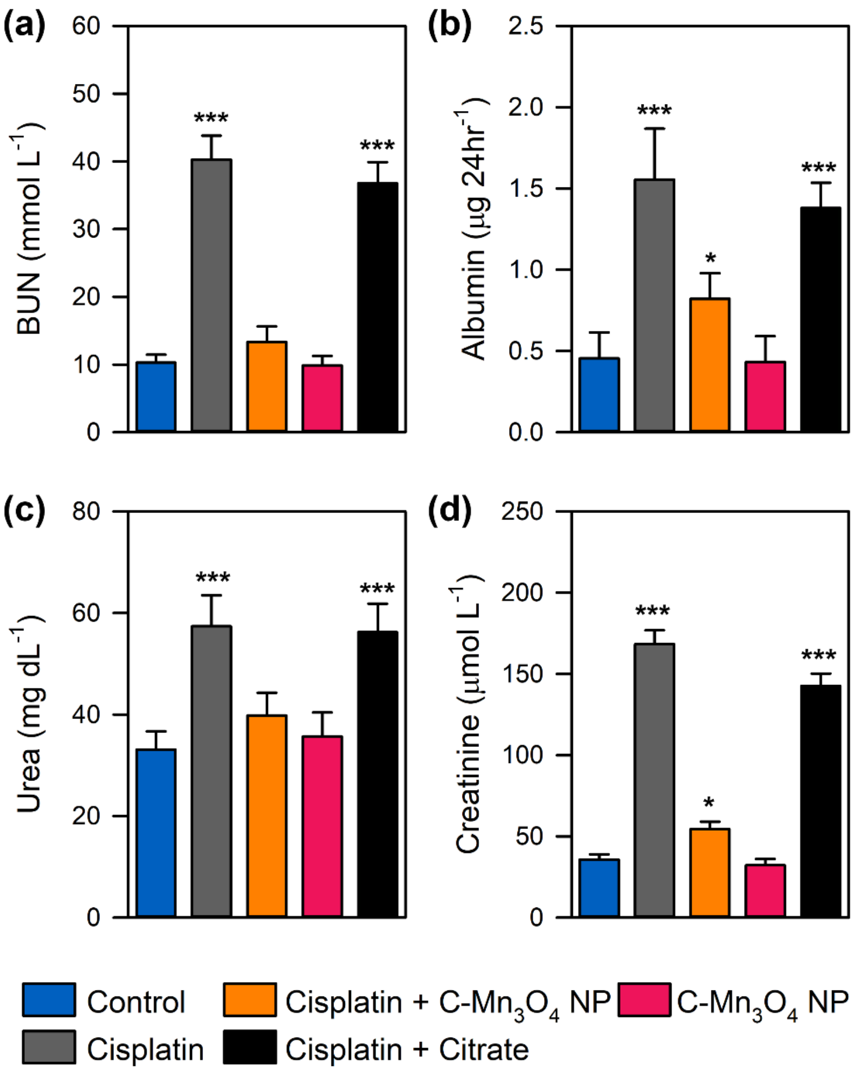


**Supplementary Figure S3. Efficacy of C-$Mn_3O_4$ NPs in reversal of CKD in animal model.** (a) Blood urea nitrogen (BUN) content. (b) Urinary albumin excretion as an indicator of albuminuria, hallmark of CKD. (c) Serum urea concentration. (d) Serum creatinine level. Note, treatment with citrate (the ligand) could not reduce the cisplatin induced nephrotoxicity. So, we left the treatment group from further downstream studies.

Data are expressed as Mean ± SD. *N*=6. *, **, *** Values differ significantly from control group (without treatment) (***p < 0.001; **p < 0.01; *p < 0.05).
